# A dual role for PGLYRP1 in host defense and immune regulation during *B. pertussis* infection

**DOI:** 10.1101/2025.09.26.678899

**Authors:** David M Rickert, Sasha Cardozo, Nicholas H Carbonetti, William E Goldman, Karen M Scanlon, Ciaran Skerry

## Abstract

*Bordetella pertussis*, the etiologic agent of whooping cough, remains a serious public health concern despite widespread vaccination. Improved therapeutics and vaccines are urgently needed to treat and prevent pertussis disease. Host recognition of bacterial peptidoglycan (PGN), including *B. pertussis* extracellular PGN fragment tracheal cytotoxin (TCT), shapes the immune response to infection. Peptidoglycan recognition proteins (PGLYRPs) are a conserved family of innate immune molecules which bind bacterial PGN. While they function as immune signaling receptors in arthropods (termed PGRPs in arthropods), PGLYRPs in mammals have thus far been primarily recognized for their bactericidal activity. Previously thought to function only as antimicrobial peptides in mammals, the immune modulatory roles of this family of peptidoglycan recognition proteins are beginning to gain greater appreciation. Peptidoglycan recognition protein 1 (PGLYRP1) is a secreted antimicrobial protein. However, its role in mammalian host defenses and immune signaling during infection with Gram-negative pathogens, such as *B. pertussis*, remain largely unknown. Here, we identify a dual role for PGLYRP1 in modulating host immune responses to *B. pertussis*. Using knockout mice, single-cell and bulk transcriptomics and functional assays, we show that PGLYRP1 contributes to host antibacterial responses to *B. pertussis*. PGLYRP1 also dampens inflammatory responses and paradoxically inhibits bacterial killing later in infection. Mechanistically, PGLYRP1 enhances nucleotide oligomerization domain (NOD)-1 signaling in response to TCT while suppressing NOD2- and triggering receptor expressed on myeloid cells-1 (TREM-1)-mediated inflammatory pathways. TCT-bound PGLYRP1 selectively impairs TREM-1 activation compared to PGNs from other bacteria, revealing a novel bacterial immune evasion strategy. These findings demonstrate that *B. pertussis* co-opts PGLYRP1 to temper inflammation and alter immune signaling, revealing a novel immune evasion mechanism of manipulating the availability and structure of their exogenous peptidoglycan, revealing implications for host-pathogen evolution, vaccine design and host-directed therapeutics.

## INTRODUCTION

*Bordetella pertussis* is a highly contagious respiratory pathogen and the causative agent of whooping cough, a disease characterized by severe, prolonged coughing and airway inflammation^1-3^. Despite widespread vaccination efforts, *B. pertussis* remains a public health concern due to waning vaccine-induced immunity and the pathogen’s capacity to evade host immune defenses^4^. Host detection of microbial components, including peptidoglycan fragments, is essential for initiating immune responses to infection. Understanding how these host-pathogen interactions shape disease outcomes is critical for developing improved strategies to fight infections.

Peptidoglycan (PGN) is a fundamental component of the bacterial cell wall, composed of repeating disaccharide chains cross-linked by short peptides^5^. PGN structure is an important determinant of innate immune recognition and responses^6, 7^. During cell wall turnover, some pathogenic bacterial species, including *B. pertussis*, release PGN fragments into the extracellular environment^8-10^. The biological roles of these fragments in infection and immunity are poorly defined.

*B. pertussis* releases a disaccharide-tetrapeptide monomeric PGN fragment known as tracheal cytotoxin (TCT)^11, 12^. TCT differs from *B. pertussis* cell wall PGN by having a 1,6-anhydro group on its MurNAc sugar due to incomplete PGN turnover^11^. The functional consequences of this 1,6-anhydro MurNAc modification are poorly understood^13^. TCT is recognized by the cytosolic pattern recognition receptor (PRR) NOD1 to activate NFkB-driven immune responses^14, 15^. *In vitro* and *ex vivo* studies suggest that recognition of TCT by NOD1 in non-ciliated mucus-secreting cells leads to the accumulation of nitric oxide within neighboring ciliated airway epithelial cells and subsequent ciliostasis and extrusion of ciliated cells^16, 17^. We recently demonstrated that TCT has broader immunomodulatory effects, skewing PGN-sensing toward NOD1 and away from NOD2, thereby dampening a stronger NOD2-dependent, pro-inflammatory cytokine production and generation of immune memory^18^. However, the broader immunological consequences of TCT release during infection, especially in the context of PGN-sensing by PRRs remain poorly defined.

Peptidoglycan Recognition Proteins (PGLYRPs) are a conserved family of soluble extracellular innate immune receptors that bind PGN^19^. In arthropods, PGRPs act as extracellular pattern recognition receptors (PRR) that trigger downstream immune pathways^20^. In mammals, they have been characterized as bactericidal proteins that target PGN on Gram-positive bacterial surfaces^21, 22^. However, recent studies have expanded their role beyond antimicrobial defense, suggesting function in regulating host immune responses. PGLYRP1 is a secreted protein stored in neutrophil granules, that is directly bactericidal against several bacterial species^23^. Recent work suggests that mammalian PGLYRPs also regulate adaptive immunity, including CD8+ T cell responses, in autoimmunity and cancer, and act as an accessory protein for the activation of inflammation amplifying receptor TREM-1 and NOD2-dependent signaling^24-26^.

TREM-1 is a PRR which amplifies inflammatory responses during infection^27^. Activation of TREM-1 by its ligands, including PGLYRP1-PGN complexes, enhances production of proinflammatory cytokines and chemokines^28^. Excessive TREM-1 activation is associated with inflammatory pathology in sepsis and other diseases^29^. Despite growing interest in PGLYRPs, their roles in host-pathogen interactions, particularly in the context of Gram-negative bacteria and secreted PGN fragments, remains poorly defined.

Here, we demonstrate that PGLYRP1 plays a dual role in host responses to *B. pertussis* infection, contributing to early bactericidal responses and fine-tuning inflammation in response to distinct PGN structures. We demonstrate that exogenous PGLYRP1 enhances NOD1 activation by TCT but failed to enhance muramyl dipeptide (MDP) activation of NOD2. Further, PGLYRP1 directly activates the pro-inflammatory receptor TREM-1, a response that is potentiated by sacculus derived PGN from Gram-positive bacteria *Staphylococcus aureus* or *Bacillus subtilis*. PGLYRP1-mediated activation of TREM-1 was not supported by *B. pertussis* extracellular PGN monomer TCT. These findings reveal that PGLYRP1 is not only contributes to anti-bacterial responses but suggests it may also be a context-specific immunomodulator whose function is shaped by the PGN it encounters. This work uncovers a previously unrecognized mechanism by which extracellular PGN and innate receptors together shape host inflammation during infection in a structure-specific manner.

## RESULTS

### Opposing roles of PGLYRP1 in early and late host defenses to *B. pertussis*

PGLYRP1 has known antibacterial activity *in vitro*, particularly against Gram-positive bacteria^23^. To investigate whether PGLYRP1 contributes to host microbial defenses against *B. pertussis*, we intranasally infected adult BALB/c wild-type (WT) and PGLYRP1-knockout (PGLYRP1 KO) mice and assessed lung bacterial burden at 4- and 7-days post infection (DPI). Surprisingly, PGLYRP1 had opposing impacts on bacterial burden at these time points (Fig. 1A&B). At 4DPI, PGYLRP1 KO mice exhibited significantly higher bacterial burdens compared to WT controls (p < 0.01), indicating that PGLYRP1 contributes to early bacterial control. However, by 7DPI PGLYRP1 KO mice had significantly lower bacterial burden compared to WT mice (p < 0.001), suggesting that PGLYRP1 impairs bacterial clearance later in infection, potentially by regulating or modulating host immune responses.

**Figure 1.**
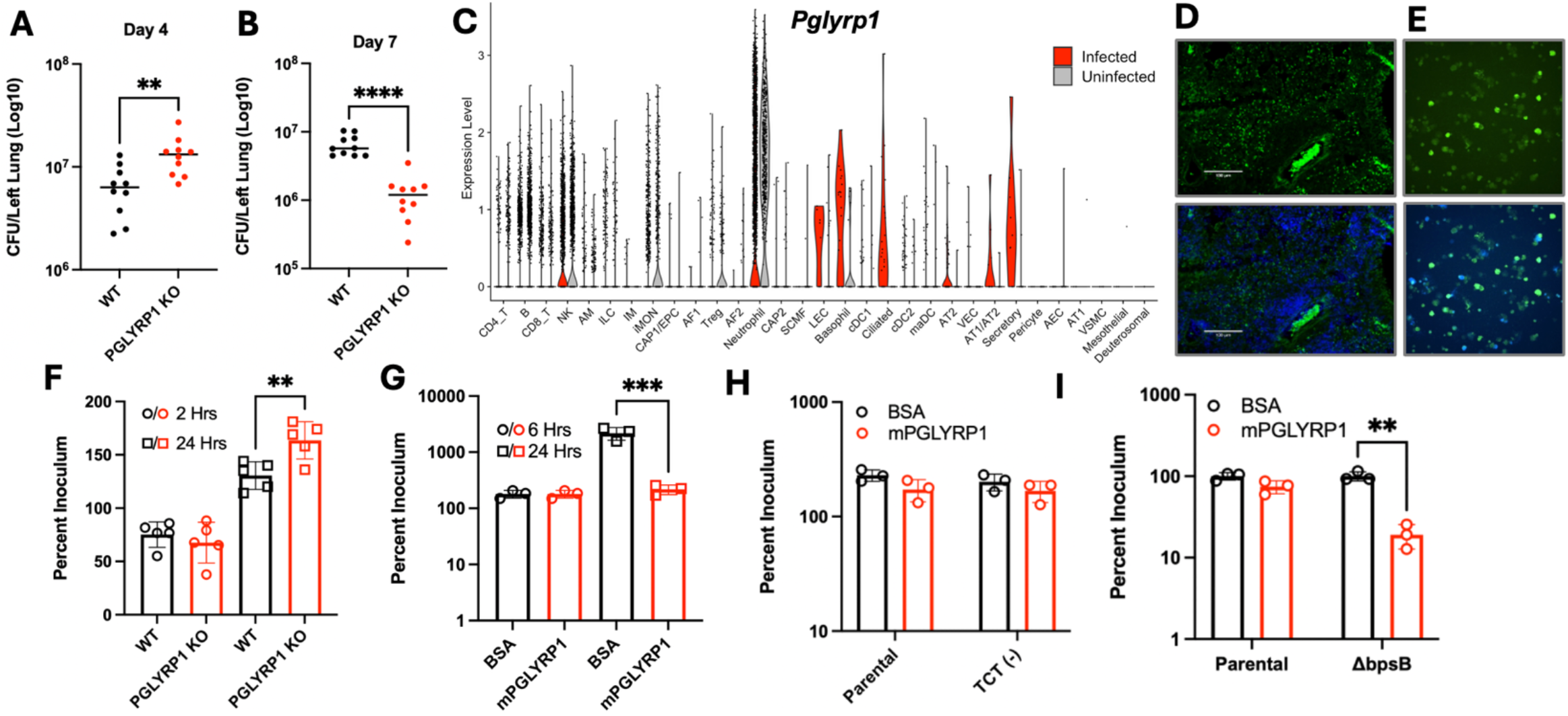
PGLYRP1 has a dual role in host defense against *Bordetella pertussis*. Lung bacterial burdens in wild-type BALB/c (WT, black) and PGLYRP1-knockout (PGLYRP1 KO, red) at 4-(A) and 7-(B) days post-infection. Violin-plot visualizing single-cell expression levels of Pglyrp1 across cell types in the lungs of C57/B6 at 4DPI for both infected and sham-challenged mice, assessed via analysis of single-cell RNA sequencing data (C). RNAscope in situ hybridization visualization of *Pglyrp1* transcripts in FFPE sections of *B. pertussis* infected mouse lung tissue (D). Each dot represents a single Pglyrp1 mRNA molecule. Images are presented at 20X and 10X magnification of *ex vivo* bone marrow derived neutrophils (E). Neutrophils were isolated from BALB/c (WT) or PGLYRP1 KO mice and incubated with *B. pertussis* for 2 (circles) or 24 (square) hours and CFU enumerated by serial dilution on Bordet Gengou agar. *In vitro* bacterial killing assay assessing recombinant murine PGLYRP1 (mPGLYRP1, red) bactericidal activity at 6- and 24-hours post-incubation compared to BSA control against WT (G) or a TCT non-releasing strain (TCT-) at 6 hours (H). Enumeration of CFU following incubation of mPGLYRP1 with wild-type *B. pertussis* (parental) or mutant lacking the *BpsB* gene of the *Bordetella* polysaccharide (Bps) operon for 6 hours (I). Data represents CFU as a percentage of starting inoculum recovered at indicated timepoint as assessed by serial dilution of culture on BG agar. Data are presented as mean ± SEM; significance determined by unpaired t-test or ANOVA as appropriate. **, *p-value* < 0.01, *** *p-value* < 0.005, **** *p-value* < 0.001

To identify which cell types express PGLYRP1 following infection, we performed single cell RNA sequencing on lungs from uninfected and *B. pertussis-*infected C57BL/6 mice at 4DPI. In uninfected lungs PGLYRP1 expression was most prominent in neutrophils along with heterogenous expression in some lymphocyte and inflammatory monocyte (iMON) populations (Fig. 1C). Upon infection, the PGLYRP1-expressing cell-types expanded to include epithelial cells like ciliated cell, secretory cells, and AT1/AT2s in addition to granulocytes like basophils (Fig. 1C). RNAscope in situ hybridization confirmed expression in the airways of infected mice and in neutrophils (Fig. 1D, E).

To determine whether PGLYRP1 contributes to antibacterial responses to *B. pertussis*, we performed *in vitro* and *ex vivo* killing assays. Based on transcriptomic data (Fig. 1C), we assessed the bactericidal potential of BMDNs from WT and PGLYRP1 KO mice incubated with *B. pertussis* for 2 or 24 hours. At 24 hours, PGLYRP1 KO neutrophils failed to control viable bacteria compared to WT neutrophils (Fig. 1F), indicating that PGLYRP1 contributes to neutrophil-mediated control of *B. pertussis*. In a cell-free assay, recombinant murine PGLYRP1 (mPGLYRP1) significantly reduced bacterial CFU after 24 hours of incubation compared to BSA control (Figure 1G, p<0.001), validating its potential antibacterial role in neutrophil mediated control of *B. pertussis* growth.

TCT release by *B. pertussis* is a product of inefficient recycling by the permease AmpG^11^. Replacing *B. pertussis* AmpG, with *E. coli* AmpG resulted in a strain (TCT-) which releases 99% less TCT (TCT(-)), while deletion of AmpG created a strain with 24-fold greater TCT release (TCT(+))^30^. Since PGLYRP1 binds bacterial PGN and the addition of excess PGN has been shown to prevent PGLYRP bactericidal activity^31^,,we hypothesized that *B. pertussis* TCT, an extracellular PGN fragment, may act as a decoy to divert PGLYRP1 activity. However, no increase in the bactericidal activity of recombinant mPGLYRP1 was noted between WT or TCT-deficient strains following incubation with PGLYRP1 (Fig. 1H), arguing against this hypothesis. Given prior work showing that *Bordetella* polysaccharides confer resistance to antimicrobial peptides^32^, we tested whether extracellular polysaccharides inhibit PGLYRP1 function. The Bps operon is required for effective production of *B. pertussis* surface polysaccharide^33^ . Using a BpsB-deficient strain we observed significantly reduced resistance to mPGLYRP1 activity compared to WT bacteria (Fig. 1I, p< 0.01), suggesting that endogenous polysaccharides can shield *B. pertussis* from PGLYRP1 activity.

Together, these results indicate that PGLYRP1 plays a dual role during *B. pertussis* infection. PGLYRP1 enhances early bacterial control via neutrophil-mediated killing and direct bacteriostatic activity but may also play a deleterious role in controlling pulmonary infection at later stages of infection (Figure 1). While TCT does not appear to interfere with PGLYRP1 activity, *Bordetella* polysaccharides, previously associated with biofilm production^33^, do, highlighting a key bacterial evasion strategy against host innate defenses.

### Peptidoglycan fragment TCT enhances PGLYRP1-mediated suppression of host responses in pertussis

Given the opposing effects of PGLYRP1 on bacterial burden at 4DPI and 7DPI, we hypothesized that PGLYRP1 exerts antibacterial activity at early time points but may suppress protective immune responses or enhance deleterious responses later in infection, impairing bacterial control by the host. Recently, PGLYRP1 has been associated with PGN recognition by innate immune receptors, including NOD2 and TREM-1^24, 26^. However, *B. pertussis* releases a NOD1 activating PGN fragment, TCT^14^.

To assess the role of PGLYRP1 in *B, pertussis*-driven inflammation, we performed histological analysis of lungs from WT and PGLYRP1 KO mice at 4 and 7DPI. We observed significantly reduced immunopathology, based on the percentage and severity of bronchovascular bundle formation and infiltration of immune cells into alveolar spaces, in PGLYRP1 KO mice at both timepoints (Fig. 2A, *p* < 0.05 at 4DPI, *p* < 0.01 at 7DPI). These findings suggest that PGLYRP1 contributes to inflammatory lung damage during *B. pertussis* infection.

**Figure 2.**
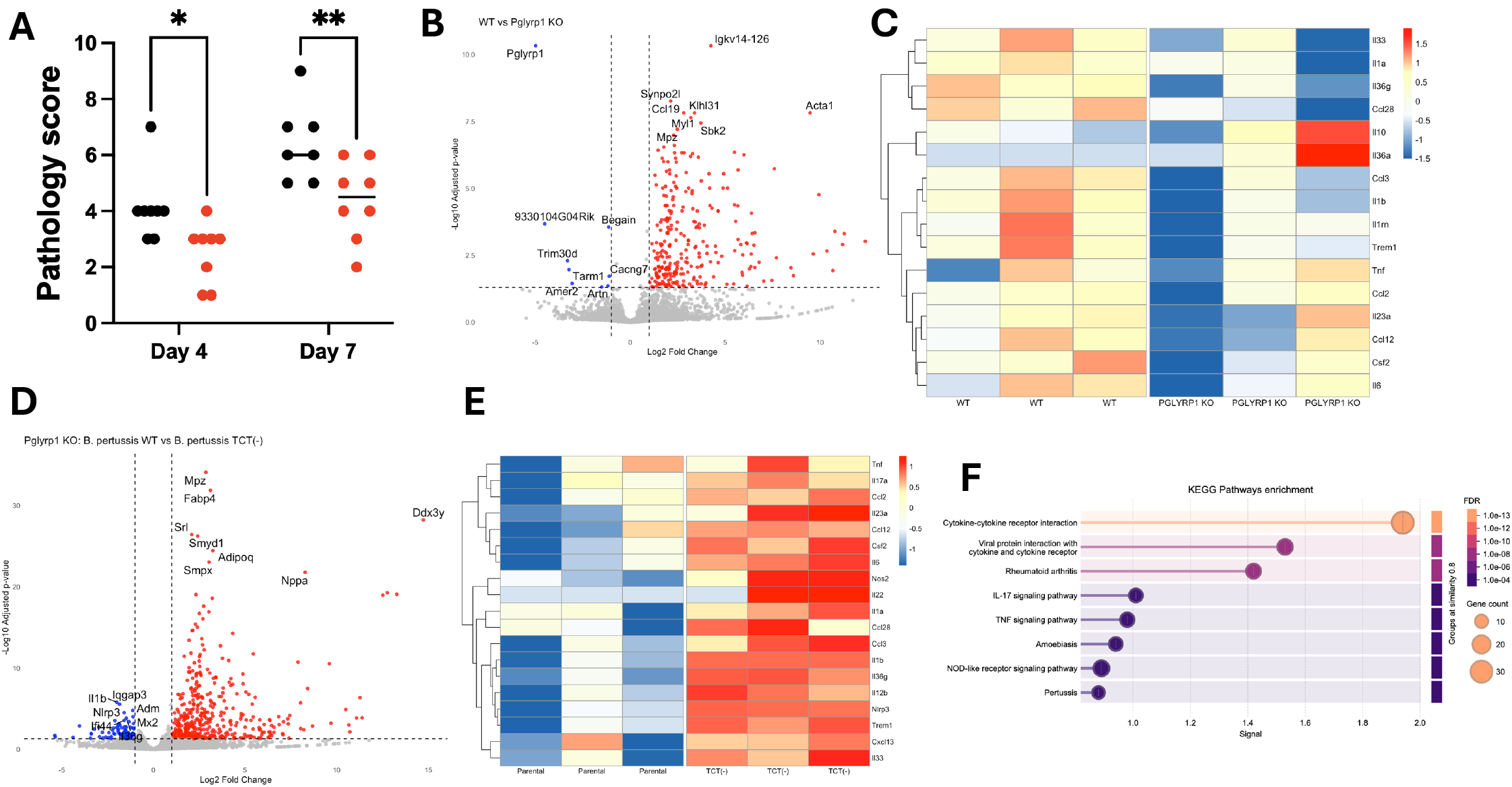
Assessment of PGLYRP1’s role in pulmonary inflammation and transcriptional responses during *Bordetella pertussis* infection. Semi-quantitative scoring of hematoxylin and eosin (H&E) stained lung sections from wild-type BALB/c (black dots) and PGLYRP1 knockout (red dots) mice at 4- and 7-days post-infection with *B. pertussis* (A). Data represents average scores of 3 blinded investigators scoring degree and percentage of peribronchial infiltration and alveolar consolidation. Bulk RNA sequencing was performed on lung homogenates collected from WT and PGLYRP1 KO mice at 4DPI to assess differential gene expression. Volcano plots (B,D) and heat-maps (C,E) showing total (B,D) and immune-related (C,E) differentially expressed genes between *B. pertussis* challenged BALB/c (WT) and PGLYRP1 KO mice (B,C) or PGLYRP1 KO mice challenged with parental wild-type *B. pertussis* (WT) or TCT-under releasing strain TCT(-). (F) KEGG enrichment pathway analysis was performed on differentially expressed genes from lung tissue isolated from mice challenged with WT or TCT(-) *B. pertussis*. Data represent the mean of biological replicates; statistical analyses were performed by Mann Whitney U test *p-values; *=0*.*003*, **, *p-value* < 0.01

To investigate PGLYRP1s role in immune signaling during *B. pertussis* infection, we performed bulk RNA sequencing on lungs from infected BALB/c (WT) and PGLYRP1 KO mice. PGLYRP1 KO mice exhibited increased expression of pro-inflammatory cytokines (e.g., *IL6, IL23A, IL1A, IL1B*, and *IL36G*) and chemokines (e.g., *CCL2, CCL3, CCL12*, and *CCL28*) alongside reduced expression of anti-inflammatory cytokine *IL10* (Fig. 2B-C). This transcriptomic profile suggests a model in which PGLYRP1 dampens key cytokine responses while paradoxically promoting immunopathology.

PGLYRP1 is a soluble PGN-binding protein. *B. pertussis* releases an extracellular PGN fragment, tracheal cytotoxin (TCT). We have recently demonstrated that TCT inhibits immune responses to *B*. pertussis^18^.We hypothesized that interactions with TCT contribute to PGLYRP1 influence on host inflammatory responses to *B. pertussis*. To test this, we compared the transcriptional responses of PGLYRP1 KO mice challenged with TCT-producing (parental) or TCT-deficient *B. pertussis* strains. Both strains induced a similar pattern of cytokine upregulation in the absence of PGLYRP1, but this increased inflammation was more robust in the absence of TCT (Fig. 2D&E). Specifically, PGLYRP1 KO mice infected with TCT(-) bacteria exhibited greater expression of IL23A, IL6, IL1B, and IL36G compared to WT infection, suggesting that TCT may potentiate the immune-dampening activity of PGLYRP1.

PGLYRP1 has previously been shown to stimulate inflammatory responses to PGN^26^. These findings indicate that the immunomodulatory function of PGLYRP1 is informed by the PGN it interacts with. TCT release may be a strategy employed by *B. pertussis* to co-opt PGLYRP1 function and suppress protective inflammation, thereby promoting pathogen persistence. Supporting this, KEGG pathway analysis on differential gene expression of 4DPI WT (parental) and TCT(-) lungs confirmed enrichment of cytokine-cytokine receptor interactions, NOD signaling, and IL-17/TNF pathways in the presence of TCT (Fig. 2F).

### TCT-PGLYRP1 interactions drive NOD1 signaling

To understand how TCT modulates PGLYRP1-dependent immune signaling, we investigated known mechanisms by which PGLYRP1 influences host PRR signaling. Recent studies suggest that intracellular PGLYRP1 can act as a co-factor in NOD2 signaling^24^ and TCT is a known NOD1 agonist. Hence, we hypothesized that PGLYRP1 may enhance NOD1 signaling in response to the muropeptide TCT, a NOD1 exclusive agonist. To test this, recombinant PGLYRP1 was incubated with TCT (1 ng/mL) before stimulation of NOD1 and NOD2 reporter cells. Consistent with previous reports, TCT robustly stimulated a NOD1 response (Fig. 3A). Notably, both murine (Fig. 3A, *p <* 0.005) and human (Fig. 3B, *p* < 0.001) PGLYRP1 significantly enhanced TCT-mediated NOD1 activation compared to TCT alone. Importantly, PGLYRP1 alone did not induce NOD1 activation (Fig. 3A), suggesting the enhanced NOD1 stimulation was not by PGLYRP1 directly activating NOD1 (Fig. 3A). Interestingly, this enhanced activation was specific to NOD1 as neither TCT nor TCT+PGLYRP1 activated NOD2 reporter cells (data not shown) since TCT is not a NOD2 agonist. On the other hand, mPGLYRP1 decreased NOD2 recognition of NOD2 agonist muramyl dipeptdide (MDP, *p* < 0.01) (Fig. 3C), suggesting exogenous PGLYRP1 boosts responses to NOD1 stimulating PGN structures like TCT but dampens responses to NOD2 stimulatory PGNs.

**Figure 3.**
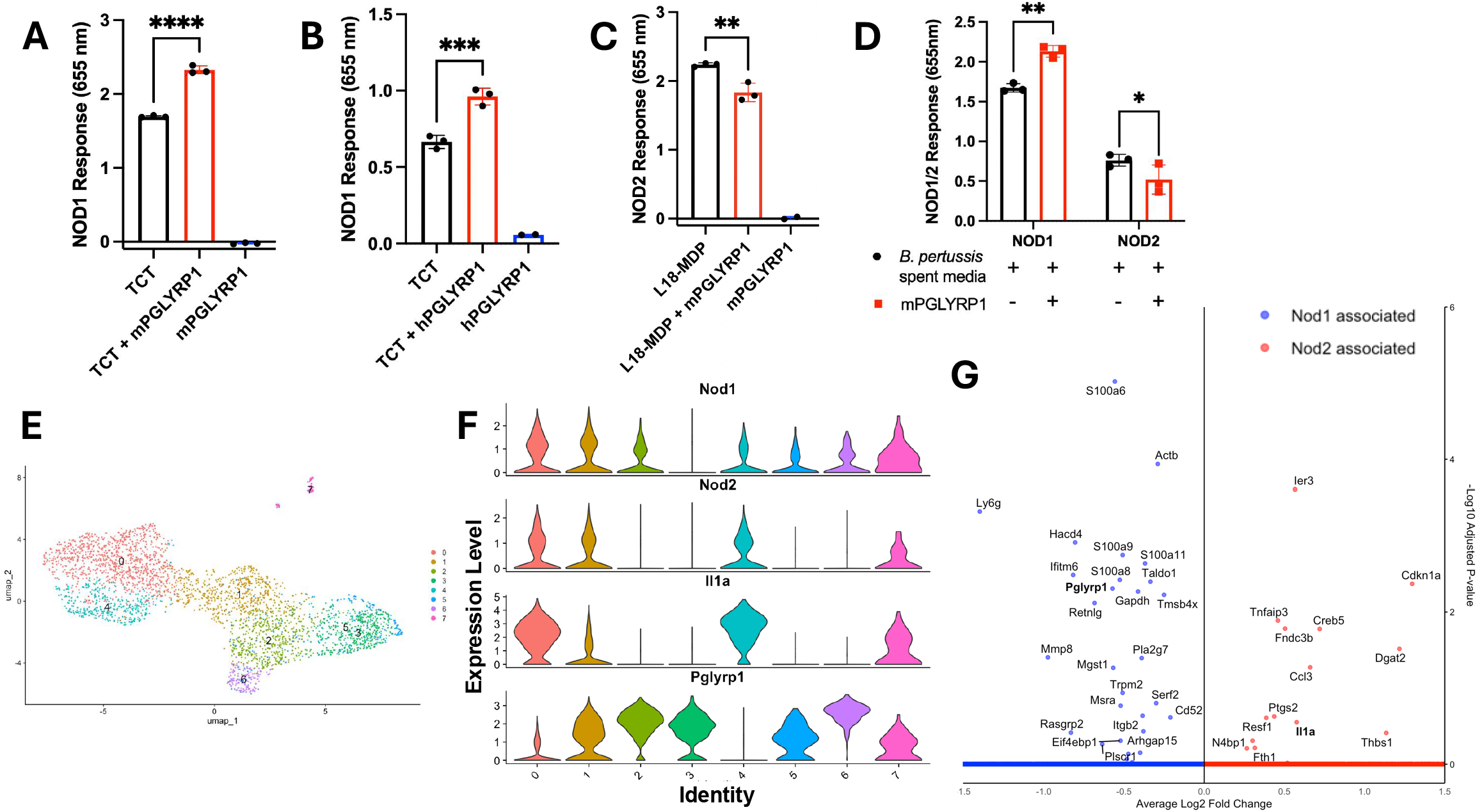
PGLYRP1 selectively modulates NOD1 and NOD2 signaling in response to *Bordetella pertussis* peptidoglycan. (A–B) HEK293 reporter cells expressing either mouse (A and D) or human (B) NOD1 or (C) NOD2 were stimulated with recombinant mouse (A) or murine or human (B-D) PGLYRP1 (12.5 ug/mL), purified tracheal cytotoxin (TCT, 1 ng/mL) (A,B), muramyl dipeptide (MDP, NOD2 agonist) (C), or conditioned bacterial growth media (OD = 0.8) (D). NF-κB-driven SEAP reporter activity was measured to assess pathway activation. Reporter activation was quantified after 18–24 hours and normalized to PBS-treated cell controls as a zero baseline. Single-cell RNA sequencing was performed on lung tissue from BALB/c mice infected with *B. pertussis* at 4 days post-infection. Neutrophils were subjected to further sub-clustering and visualized as a UMAP (E). Violin plots visualizing expression of Nod1, Nod2, IL1a and Pglyrp1 across neutrophil sub-clusters (F). Volcano plot demonstrating differentially expressed genes in Nod1 (blue) or Nod2 (red) expressing neutrophils (G) Single-cell analyses represent combined data from two mice per group. Statistical analyses were performed using Student’s t-test or adjusted p-values as appropriate. **, *p-value* < 0.01, *** *p-value* < 0.005, **** *p-value* < 0.001

To determine if PGLYRP1 modulates NOD responses to other potential PGNs released by *B. pertussis*, we stimulated reporter cells with mPGLYRP1 incubated with conditioned media from WT or TCT(-), or TCT(+) cultures. Consistent with our results using purified TCT, mPGLYRP1 amplified NOD1 signaling in response to *B. pertussis* conditioned media, likely due to increased recognition of extracellular TCT (Fig. 3D, *p <* 0.01). In contrast, NOD2 signaling was significantly suppressed by mPGLYRP1 (Fig. 3D), consistent with our results with purified PGN (Fig. 3A-C). Together, these findings suggest that exogenous PGLYRP1 acts as a selective modulator of NOD signaling, enhancing NOD1 activation while dampening NOD2 responses.

To assess the functional consequence of skewed NOD1 versus NOD2 expression and signaling during infection, we analyzed single-cell RNA sequencing data from lungs of BALB/c mice at 4DPI. We examined NOD1 and NOD2 expression across neutrophil sub-clusters (Fig. 3E&F), as neutrophils are the primary expressors of PGLYRP1 during *B. pertussis* infection (Fig. 1C). NOD2-expressing neutrophils showed higher expression of pro-inflammatory genes including *Il1a, Ccl3* and *Ptgs2* (COX-2) (Fig. 3F&G). In contrast, NOD1 expression correlated with genes involved in anti-inflammatory immune responses: *Pla2g7* (degrades platelet activating factor), *Tmsb4x* which inhibits neutrophil migration, promotes repair and inhibits NFkB and increased expression of PGLYRP1, which may further positively regulate these pathways^34^. This identifies a correlation between pro-inflammatory cytokine expression and NOD2 signaling in neutrophils.

Interestingly, when we further analyzed the clusters with high Nod2 expression, like clusters 0 and 4, we observed gene expression patterns which correspond to previously identified inflammatory neutrophil populations. Cluster 0 hallmark genes were associated with interferon-stimulated programming (*Calhm6, Cxcl9, Tma16, Slamf7, Baft2*) (Sup. Fig. 1), and *B. pertussis* infection has been shown to cause both deleterious and beneficial interferon responses^34-36^. Additionally, poor *B. pertussis* host-responses have been linked to interferon-stimulated neutrophils, providing interesting context to this identified neutrophil cluster^37, 38^. Furthermore, cluster 4 marker (*Hilpda, Csf3, Myl4, Odc1, Spp1*) neutrophils have been identified in tumor exhaustion environments, and related to hypoxia response, highlighting another inflammatory neutrophil population (Sup. Fig. 1)^39^.

These results highlight the heightened inflammatory phenotype of NOD2+ neutrophils and suggest that TCT-mediated skewing of NOD responses towards NOD1, amplified by PGLYRP1, may be a mechanism to suppress inflammation and promote *B. pertussis* persistence.

### TCT-PGLYRP1 complexes dampen TREM-1 activation, limiting inflammation during *B. pertussis* infection

In addition to their role in NOD signaling, PGLYRP1-PGN complexes have been shown to activate TREM-1, driving NFkB-mediated inflammatory responses^26^. This ability to activate TREM-1 is boosted by the presence of a PGN from the sacculi of *S. aureus, B. subtilis or E. coli*^26^. To determine if TREM-1 contributes to host responses to *B. pertussis*, we first assessed TREM-1 expressing cell-types following *B. pertussis* infection in mice using single cell RNA-sequencing. Following *B. pertussis* infection, TREM-1 was primarily expressed by neutrophils, with lesser expression noted in monocytes and macrophages (Fig. 4A).

**Figure 4.**
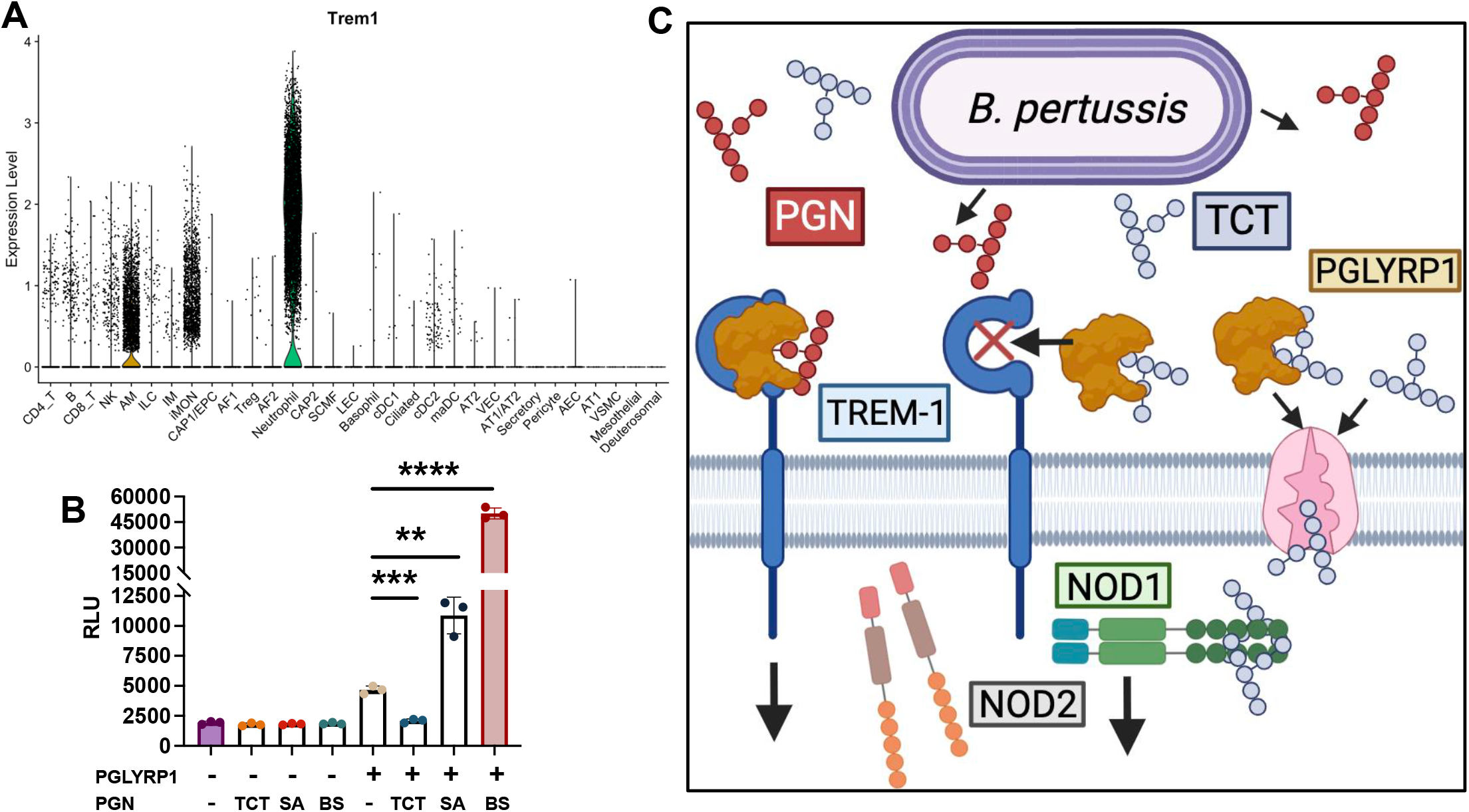
TCT inhibits PGLYRP1 signaling via TREM-1 Violin plot generated from single-cell RNA sequencing (scRNA-seq) displaying TREM-1 expression on neutrophils (green), alveolar macrophages (AM, gold), and inflammatory monocytes (IM) at 4DP1 (A). TREM-1 activation measured via NFAT-driven luminescence (relative luminescence units, RLU) in TREM-1 reporter cells treated with recombinant human PGLYRP1 with or without 5µg/mL of Lys-type peptidoglycan derived from *Staphylococcus aureus* (5 µg/mL), TCT (5 µg/mL) or a DAP-type PGN from Bacillus subtilis (5 µg/mL) (B). Diagram depicting proposed model (C). All TREM-1 activation assays were performed with biological triplicates (n = 3). Statistical analyses were performed using unpaired two-tailed Student’s t-test. *, *p-value <* 0.05, **, *p-value* < 0.01, *** *p-value* < 0.005, **** *p-value* < 0.001

To determine if TCT influences PGLYRP1-dependent TREM-1 activation, we assessed the ability of TCT, a DAP-type PGN monomer, to enhance PGLYRP1-mediated TREM-1 activation. As reported previously, preparations of peptidoglycans isolated from *S. aureus* and *B. subtilis* enhanced PGLYRP1-mediated activation of a TREM-1 reporter cell line (Fig. 4B, *p* < 0.01)^40, 41^. Given the anti-inflammatory role of TCT observed in our data (^18^ and Figs 2 and 3), and known role for TREM-1 in *B. pertussis* pathogenesis^42^ we proposed that TCT may be a poorer enhancer of PGLYRP-1 mediated TREM-1 activation. Consistent with our hypothesis the addition of TCT significantly reduced TREM-1 activity (Fig. 4C). This suggests that TCT, unlike PGN from the sacculi, fails to support and possibly even antagonizes PGLYRP1’s ability to activate TREM-1 (Fig. 4C). Further studies are required to discern the contribution of PGN structure and solubility in this model.. Combined with our previous work^42^, these findings highlight the importance of PGN in bacterial regulation of TREM-1 and its role in *B. pertussis* pathogenesis and disease and the potential impact of skewing TREM-1 responses.

## DISCUSSION

Our study reveals a previously unrecognized role for the PRR, PGLYRP1, in orchestrating both antibacterial defenses and inflammatory responses during *Bordetella pertussis* infection. While traditionally characterized as a bactericidal effector which mediates its effects by binding PGN on the surface of Gram-positive bacteria, emerging work suggests its role may extend to mammalian immune signaling^25^. The contribution of PGLYRP1 to PRR mediated inflammatory responses in mammals and in particular how PGN structure impacts the outcomes of these interactions, remains largely unexplored. Our findings demonstrate that PGLYRP1 can functions as a PGN structure-specific interpreter which translates structure into context and pathogen specific immune signaling. This dual role expands our understanding of how innate immune receptors fine-tune host-pathogen interactions in the airway based on the structure of PGN encountered. Prior studies have also shown that endogenous PGLYRP1 bolstered NOD2-driven macrophage activation in response to GMTriP-K, but not in response to related PGN fragments such as GMDiP or MDP, highlighting the structural specificity of PGLYRP1-mediated immune recognition^24^. We expand on these findings to suggest exogenous PGLYRP1 biases NOD1 responses and dampens NOD2 responses. Furthermore, we suggest pathogens like *B. pertussis* have evolved to release PGN structures, such as TCT, which dampen the ability of PGLYRP1 to signal via TREM-1 (Fig. 4C), suggesting an active area of ongoing host-pathogen evolution.

Here, we propose PGLYRP1 as a PGN-structure specific regulator of inflammatory responses. This is comparable to SIGLECs, which can distinguish self from non-self glycans to ignore or elicit immune responses^43^. During *B. pertussis* infection, PGLYRP1, predominantly expressed by neutrophils, contributes to early bacterial clearance, as PGLYRP1 KO mice exhibit significantly higher bacterial loads at 4DPI. The bactericidal activity of PGLYRP1 is diminished by the presence of extracellular polysaccharide, highlighting the potential for *Bordetella pertussis*-derived extracellular decoys to prevent antibiotic activity. PGLYRP1 KO mice exhibit elevated inflammatory cytokine expression, suggesting PGLYRP1 also serves an anti-inflammatory function. Transcriptomic analysis of PGLYRP1 KO lungs infected with mutants which fail to release TCT or the parental strain support the hypothesis that PGLYRP1 limits inflammation in a PGN structure-dependent manner: specifically, the monomeric PGN fragment TCT potentiates the anti-inflammatory activity of PGLYRP1. An alternative explanation for the discrepancy observed in bacterial burden at early and later timepoints of infection is that PGLYRP1-directed lysis of *B. pertussis* releases more host-available muropeptides, which PGLYRP1 can then interact with to enhance less protective NOD1-driven immune responses at the expense of more potent NOD2-driven responses (Fig 1)^18^.

Mechanistically, PGLYRP1 enhances NOD1 signaling in response to TCT while suppressing NOD2 signaling in response to MDP directing the polarization of immune responses away from a more inflammatory NOD2-driven response. Single cell RNA sequencing revealed that PGLYRP1 expression is enriched in NOD1+ neutrophils, which have less inflammatory profiles than NOD2+ neutrophils. These data suggest *B. pertussis* may exploit PGLYRP1 to amplify a NOD1-dominant signaling environment that blunts host inflammation. Recent work from our group highlighted the ability of TCT to polarize NOD responses, towards NOD1 as an immune evasion mechanism^18^. This work expands on this finding, highlighting the role of PGLYRP1 and PGN structure in these findings, but further work should validate and expand on how PGLYRP1 modulates responses to muropeptide outside the cellular and murine models used in these studies.

These findings shift our understanding of mammalian PGLYRP1 from a passive bactericidal protein to an immunomodulator, whose contribution to inflammation is determined by the structure of PGN it encounters. Additionally, they inform our views of PGN structure and how it impacts host responses, suggesting bacterial release of PGNs, as a mechanism to dampen host immunity. These results propose a novel paradigm in host-pathogen interactions; that bacteria may release structurally tailored PGN fragments to bias host immune sensors and signaling in their favor.

Excitingly, we report a novel axis by which PGLYRP1 regulates TREM-1, a known amplifier of inflammation. TREM-1 activation has been reported to strengthen NF-kB signaling pathways which NLRs like NOD1 and NOD2 canonically signal through^27^. While PGLYRP1 synergized with PGNs from *S. aureus* and a *B. subtilis* to enhance TREM-1 activation, the presence of TCT antagonized this effect (Fig. 4). There are several differences in the structures of these PGNs: unlike monomeric TCT, PGN tested from other bacteria are derived from the sacculus and are insoluble, additionally *B. subtilis* PGN is amidated. Further studies are required to discern if solubility, size or other factors are responsible for differences in the effect on PGLYRP1-mediated TREM-1 activation. Additionally, TCT contains a 1,6 anhydroMurNAc. Bifidobacterium have been predicted to produce abundant anhydro-PGNs which markedly reduce LPS-stimulated cytokine expression in RAW264.7 cells, underscoring a potential anti-inflammatory effect of 1,6-anhydro-MurNAc containing PGNs, such as TCT.

Previous work from our group demonstrated that inhibition of TREM-1 prevented immunopathology and inhibited cytokine and chemokine responses to infection, highlighting the potential utility of disturbed TREM-1 signaling for *B. pertussis*^*42*^. TREM-1 inhibitor treated mice showed reduced inflammation and inflammatory cytokine responses following *B. pertussis* infection highlighting the importance of this pathway in pertussis pathogenesis and the potential utility of its manipulation by bacteria. Further, our findings suggest that selectively targeting TREM-1 or PGN structure sensing by PGLYRP1 could attenuate disease severity without compromising bacterial clearance, a strategy aligned with emerging host-directed therapeutics in TB and sepsis^44-46^. However, the role of these players in generating effective antibody responses still needs to be elucidated so that we can understand the implications on long-term immunity.

We hypothesize that *B. pertussis* may attenuate host immune responses not merely by evading recognition, but by actively hijacking host regulatory circuits through the release of specific PGN structures. Taken together, our findings support a model in which PGLYRP1 serves dual and temporally distinct roles during infection, by promoting bacterial killing while restraining inflammation in a PGN-structure specific manner. This duality may reflect temporal or disease stage specific changes in the types of PGN available to host PRRs, as 1,6-anhydro MurNAc containing PGNs are usually internal, unless released following bacterial lysis. The ability of *B. pertussis* to exploit this shift by releasing TCT to suppress excessive inflammation represents a novel immune evasion strategy.

More broadly, this work demonstrates that the structure of PGN fragments is a key determinant of host immune outcomes. Innate receptors like PGLYRP1 relay this structural information to shape both defense and pathology, providing an avenue for pathogen hijacking by alternating the availability of different PGN structures. Targeting the TCT–PGLYRP1–TREM-1 axis may offer a novel therapeutic or vaccine-mediated approach to mitigate immunopathology in pertussis and potentially other infections involving secreted PGN. These findings reframe PGN as both a pathogen-associated molecular pattern and an immunomodulatory cue that both host and pathogen exploit to regulate the immune-inflammatory environment. Future studies using defined synthetic muropeptides will be necessary to pinpoint structural determinants of the reduced immune activation noted toward TCT. Likewise, how these alterations in PGN structure impact PGLYRP1 interactions with host PRRs remains unclear.

## METHODS AND MATERIALS

### Mice and Ethics Statement

All animal procedures were approved by the Institutional Animal Care and Use Committee at the University of Maryland School of Medicine (protocol 00000108). Adult (6–8-week-old) BALB/c (Charles River, 028) and PGLYRP1 knockout (Pglyrp1^tm1Rdz^/Pglyrp1^tm1Rdz^ in BALB/c background, generously provided by Dr. Roman Dziarski^47^), mice were bred and housed under specific pathogen-free conditions. To control for and assess the potential of sex-related differences in pertussis pathogenesis both male and female mice were used in experiments. For transcriptomic studies, RNA was isolated from the lungs of PBS- or *B. pertussis* challenged female mice.

### Bacterial Strains and Infections

*B. pertussis* strains, including wild-type BC36, a streptomycin-resistant Tohama I derivative, or isogenic mutants (TCT-, TCT+, BpsB KO) were cultured on Bordet-Gengou agar supplemented with 10% defibrinated sheep blood for in vivo infections or in Stainer-Scholte (SS) broth for in vitro assays. Strains were grown in SS broth to mid-log stage (OD = 0.8), and the resulting media was centrifuged and filtered to remove any bacteria to make the conditioned media used for cell culture assays. TCT mutants were generously supplied by William Goldman (University of North Carolina at Chapel Hill). TCT-strain was generated by expression of the *E. coli* AmpG in BC36, resulting in a 50-fold decrease in TCT release. TCT+ was generated by replacing native *B. pertussis* AmpG with a kanamycin resistance cassette, this resulted in a 24-fold increase in TCT release^30^. Adult mice were anesthetized and inoculated intranasally with 2 × 10^6^ CFU of *B. pertussis* in 50 µL PBS. Lungs were harvested at 4DPI and 7DPI for bacterial burden analysis by plating of serially-diluted lung homogenates.

### RNA isolation and processing

Lung tissue was harvested at 2 or 7DPI for RNA isolation by the Trizol-choloroform method. Briefly, lung tissue was homogenized using the Omni Bead Beader (Omni, Inc), phase separated using chloroform, and precipitated using isopropanol. RNA was then quantified and converted to cDNA (iScript cDNA Synthesis Kit). Quantitative real-time PCR (qPCR) was performed using the BIORAD CFX96 real-time PCR instrument. Gene expression was calculated as fold change relative to PBS-inoculated control animals using the 2^^^−ΔΔCT threshold cycle method and normalized to Hypoxanthine phosphoribosyltransferase (HPRT) (internal housekeeping gene).

### Bulk RNA Sequencing

At 4DPI, lungs were harvested from 3 WT or PGLYRP1 KO BALB/c mice infected with WT or TCT-deficient *B. pertussis*. Total RNA was extracted from homogenized lungs using Trizol Reagent (Invitrogen) according to the manufacturer’s protocol. RNA integrity was confirmed using a Bioanalyzer (Agilent), alignments were generated by HISAT2 sequencing performed using a Illumina NextSeq 1000. FastQC and RSeQC were used to assess read and alignment quality. Samtools was used to generate alignment statistics. Reads were aligned with the *Mus musculus* GRCm39 reference genome. Differential gene expression was analyzed using DESeq2, with significance defined as adjusted p-value < 0.05.

### Single-Cell RNA Sequencing

Single-cell suspensions were prepared from lung tissues of duplicate infected and uninfected control C57BL/6 mice at 4DPI, barcoded and sequenced as individual mice. Following red blood cell lysis, cell viability was assessed by trypan blue staining, viable cells were sorted and loaded onto the 10x Genomics Chromium Controller for barcoding and library construction. Sequencing was performed on an Illumina NovaSeq platform. Data were processed using Cell Ranger and analyzed using the Seurat package in R. Quality control cutoffs of between 1000 and 10000 features and <20% mitochondrial DNA. Data were normalized and scaled in Seurat v5, and groups were integrated via Harmony-based integration. Clusters were identified, determined, and visualized via Seurat library workflow (RunPCA, FindNeighbors, FindClusters, RunUMAP) and then annotated via hallmark markers determined by differential gene expression analysis across clusters using Wilcoxon rank-sum tests. This dataset is now deposited with the Gene Expression Omnibus (GSE324217)

### Neutrophil Isolation and Bactericidal Assays

Bone marrow-derived neutrophils (BMDNs) were isolated by density gradient centrifugation from adult C57BL6 mice and cultured in DMEM supplemented with 10% FBS. For killing assays, BMDNs were incubated with *B. pertussis* at a multiplicity of infection (MOI) of 10:1 for 2 or 24 hours. Supernatants were serially diluted and plated to quantify surviving CFU. Recombinant murine PGLYRP1 (R&D Systems) or BSA control was incubated with live *B. pertussis* or purified PGN fragments (TCT gifted by Prof. William Goldman or *S. aureus* PGN, Invivogen) to assess direct bactericidal activity.

### RNAscope In Situ Hybridization

RNA in situ hybridization was performed on 5um formalin fixed paraffin embedded lung sections with the RNAscope detection kit to confirm RNA sequencing identification of PGLYRP1 transcripts (Advanced Cell Diagnostics). Hybridization, amplification, and detection steps were carried out according to the manufacturer’s protocol. Briefly, slides were deparaffinized in xylene and rehydrated using an ethanol series. Sections were treated with hydrogen peroxide for 10 minutes at room temperature, followed by treatment with target retrieval reagent (Advanced Cell Diagnostics). Sections were hybridized with target-specific probes (Advanced Cell Diagnostics) followed by amplification and fluorescent detection (Opal 570 and Opal 520). Slides were counterstained with hematoxylin and eosin and imaged using an ECHO Revolve 4 and analyzed with ImageJ.

### Reporter Cell Assays for NOD1, NOD2, and TREM-1 Activation

HEK293-derived reporter cells expressing murine or human NOD1 and NOD2 with an, or TREM-1 and an NFkB–dependent luciferase or NF-kB inducible SEAP reporter (InvivoGen) and TREM-1-DAP12 reporter cells (Eurofins Discover X )in the Jurkat T cell background with an NFAT-dependent luciferase were stimulated with *B. pertussis* tracheal cytotoxin TCT (1 ng/mL), conditioned media (1:20 ratio of conditioned bacterial media to cell culture media), PGN fragments (TCT, *S. aureus*, or *B. subtilis* PGN (Invivogen)) in the presence or absence of recombinant human PGLYRP1 (R&D systems) at a concentration of 5ug per well for 24 hours. Supernatants were analyzed for reporter activity using colorimetric (NOD1 and NOD2 reporter cells; Promega) or luminescent (TREM-1 reporter cells; Invivogen) detection kits (Promega). TCT was quantitated from its peak area elution profile from the final purification step, For each preparation of TCT this area was compared to the peak area of a purified standard that had been quantitated by amino acid analysis and generously provided by William Goldman, (UNC)

### Histopathology

Lungs were fixed in 10% formalin, embedded in paraffin, and sectioned at 5 µm thickness. Slides were stained with hematoxylin and eosin (H&E). Scores are given as the average score from 3 blinded investigators based on percentage and severity of bronchovascular bundle formation and infiltration of immune cells into alveolar spaces.

## Statistics

Significant differences in pathology scores were calculated at each timepoint using a 2-tailed Mann-Whitney U test. Each timepoint was analyzed independently. A p-value < 0.05 was considered significant. A p-value < 0.05 is denoted by *, < 0.01 by ** and < 0.005 by ***

## ACKNOWLEDGEMENTS

This work was supported by NIH award R01AI167947. Prof. Roman Dziarski (Indiana University School of Medicine) generously provided PGLYRP1 KO mice. Prof. Rajendar Deora (Ohio State College of Medicine) generously provided the BpsB deletion mutant strain of *B. pertussis*.

## Notes

https://www.ncbi.nlm.nih.gov/geo/query/acc.cgi?acc=GSE324217

### Competing Interest Statement

The authors have declared no competing interest.

### Summary of Updates

Text updated following reviewer critiques. Language improved throughout manuscript. Figure 1 edited to show uninfected and infected datasets.

